# Neuroimaging-based Individualized Prediction of Cognition and Behavior for Mental Disorders and Health: Methods and Promises

**DOI:** 10.1101/2020.02.22.961136

**Authors:** Jing Sui, Rongtao Jiang, Juan Bustillo, Vince Calhoun

**Author notes:** **Address for Correspondence:** Vince Calhoun Ph.D. and Jing Sui Ph.D., Tri-institutional Center for Translational Research in Neuroimaging and Data Science (TReNDS), Georgia State University, Georgia Institute of Technology, and Emory University, Atlanta, GA, USA, 30303. Jing Sui and Rongtao Jiang contribute equally as co-first authors.

## Abstract

The neuroimaging community has witnessed a paradigm shift in biomarker discovery from using traditional univariate brain mapping approaches to multivariate predictive models, allowing the field to move towards a translational neuroscience era. Regression-based multivariate models (hereafter “predictive modeling”) provide a powerful and widely-used approach to predict human behavior with neuroimaging features. These studies maintain a focus on decoding individual differences in a continuously behavioral phenotype from neuroimaging data, opening up an exciting opportunity to describe the human brain at the single-subject level. In this survey, we provide an overview of recent studies that utilize machine learning approaches to identify neuroimaging predictors over the past decade. We first review regression-based approaches and highlight connectome-based predictive modeling (CPM), which has grown in popularity in recent years. Next, we systematically describe recent representative studies using these tools in the context of cognitive function, symptom severity, personality traits and emotion processing. Finally, we highlight a few challenges related to combining multimodal data, longitudinal prediction, external validations and the employment of deep learning methods that have emerged from our review of the existing literature, as well as present some promising and challenging future directions.

## Introduction

A biomarker, or biological marker, generally refers to a measurable indicator of normal biological processes, pathogenic processes, or pharmacologic responses to a therapeutic intervention (1). The ultimate goal of developing biomarkers is to perform individual-level predictions of human behavior that may eventually benefit educational or clinical practices in real-world settings (2). However, most neuroimaging findings from the published studies cannot be easily translated into tools with practical utility. On one hand, many existing studies primarily focus on group-level mapping using univariate analytical techniques across a massive number of isolated brain measurements, e.g., either detecting brain features showing significant group differences between healthy controls and patients with brain disorders using statistical inferences, or establishing the brain-behavior relationship using correlational analysis. Despite the fact that these investigations have offered valuable insights into the human brain, such studies may dilute the considerable heterogeneity within group that is crucial for understanding the related neurobiological basis (3). On the other hand, due in part to a lack of out-of-sample validation, group-level inferences often overfit to both the signal and noise in a specific dataset. Therefore, the generalizability of such group-level findings to new individuals remains unknown (4). In addition, traditional univariate research focuses on explaining the neural correlates of observed behavior, rather than predicting future behavior based on neuro(5)imaging signatures, the latter being an important aspect of moving towards a translational neuroscience era (6).

Recent years have witnessed a paradigm shift in biomarker discovery from using traditional univariate brain mapping techniques to multivariate predictive models for an individual (7). Different from conventional approaches, machine learning-based methods can establish integrated brain models by taking into account the multivariate nature of diverse brain function or structure measurements across the whole brain (8), which may open up an exciting opportunity to analyze neuroimaging data at the single-subject level (9). Hence, the application of machine learning approaches that can facilitate the search for reliable neuroimaging biomarkers in both health and disease has been a highly discussed topic (5).

While most predictive analyses with neuroimaging data focus on dichotomous classification, there is an emerging trend in using regression-based machine learning approaches to reveal individual differences in disease severity or cognitive functioning from brain imaging data, *i.e.*, the individualized prediction of human behaviors (3). Compared to binary classification, behavior prediction can be more challenging, since it considers the problem of quantitatively estimating the specific scores for a continuous behavioral measure over the whole range of the variable, instead of determining the group membership (10). Nevertheless, such applications can tell us more about the severity of symptom for a psychotic patient, the level of negative affect an individual tends to experience, or how well a participant can perform during a cognitive task. The biggest difference between predictive modeling and conventional correlational analysis is that predictive modeling generally employs a built-in cross-validation strategy to guard against the possibility of overfitting, which holds substantial promise for testing the generalizability of the identified biomarkers (11). The prediction of phenotypic measures requires dedicated design and techniques, recent reviews provide practical guidance on this topics (11, 12).

In this review, we will focus on approaches and cases specifically relevant to “cognitive biomarker” identification. We first outline various machine learning approaches and some key aspects on regression-based prediction, which aims to decode individual differences in a continuous behavioral phenotype from neuroimaging data. Next, we review studies on predictive modeling identified via a key word search in PubMed and Google Scholar over the past decade. Finally, challenges and future directions in the field of predictive modeling are discussed.

### Study search criteria and results

Over the past few years, there has been increasing interest in using regression-based predictive modeling to determining cognition-related biomarkers. To summarize these studies, we performed a systematic review to include papers in English based on a search on PubMed and Google Scholar using search terms: “biomarker”, “cogniti*”, “predict*”, “behavior*”, “regression”, “individual difference”, “neuroimaging”, “machine learning”, and “cross validation”, both in isolation and in combination. Searches were restricted to journal papers from January 2010 to October 2019. More than 500 papers were identified. We further reviewed the abstract and the full-text to restrict the papers to predictive studies that utilized regression-based machine learning approaches, employed cross-validation or external validation strategy to assess model generalizability, and reported the prediction accuracy using correlation coefficient. These criteria led to the inclusion of 122 papers on the topic of behavior prediction. Note that many studies performed prediction for more than one behavioral metric or several sub-dimensions of one cognitive scale. Overall, a total of 340 metric prediction accuracies were reported by all these 122 studies. **Table S1** lists these studies in terms of employed imaging modality, behavioral measure of interest, sample size, adopted method, and prediction accuracy. We summarize these papers and illustrate some key aspects in **Figure 1**.

**Figure 1.**
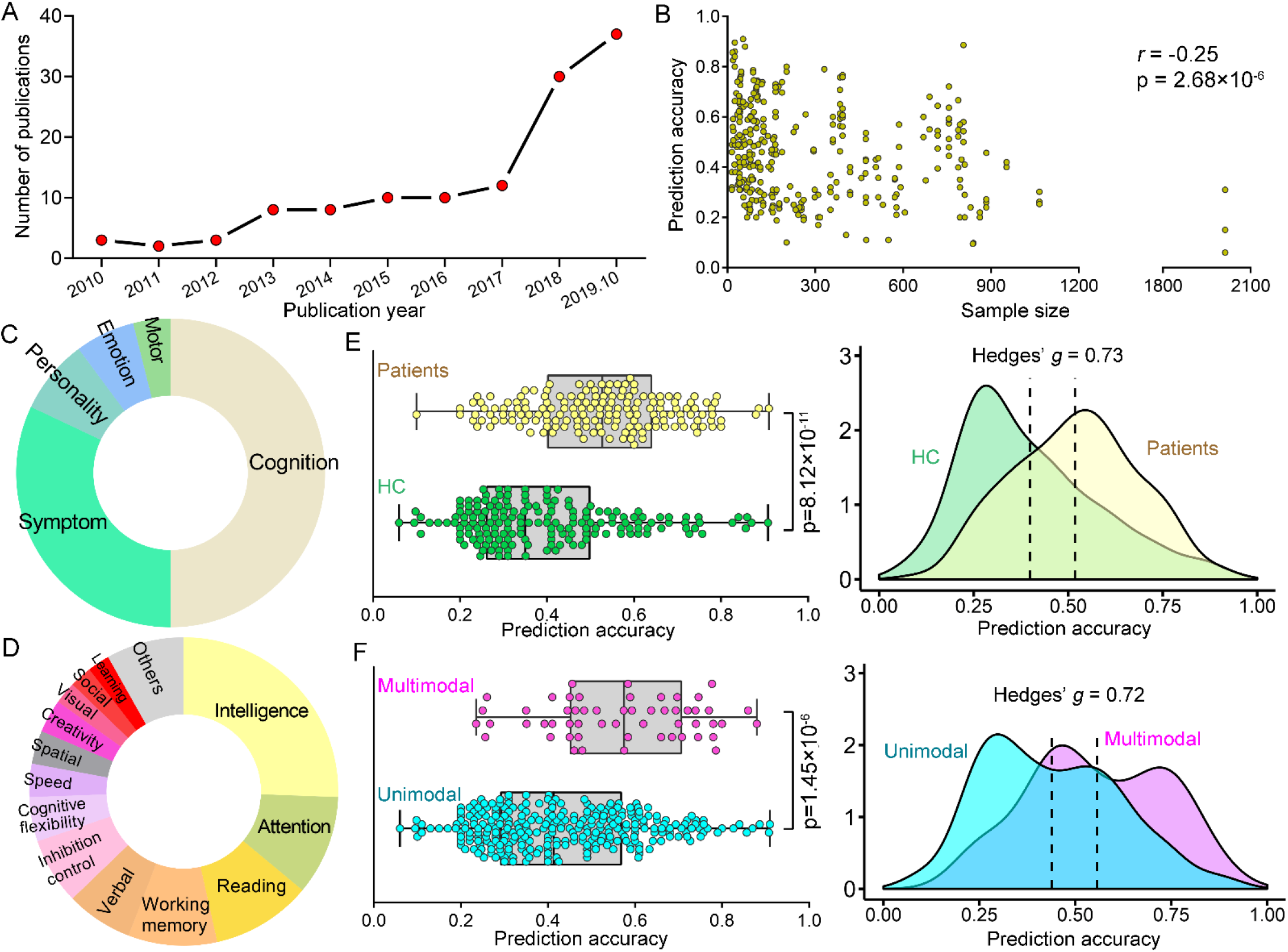
Visual summary of studies using regression-based machine learning approaches to predict continuous variables. (**A**) There is an obviously increasing trend in the number of papers published each year since 2010. (**B**) The overall prediction accuracy against the corresponding sample size used in the studies. (**C**) The type of behaviors of interest that are used as the target measures among all surveyed studies. (**D**) Cognitive metrics adopted in our surveyed studies. (**E**) Distribution of prediction accuracy for healthy subjects and patients with brain disorders shown as boxplot plot (left) and kernel density (right). (**F**) Distribution of prediction accuracy from studies using multimodal or unimodal data.

### Sample size vs Prediction accuracy

There is an obviously increasing trend in the number of papers published each year (**Figure 1A**). Most of these studies have fewer than 200 participants, and a sample size of about 100 is the most common case (**Figure 1B**). In general, prediction accuracies exhibit a significantly negative correlation (*r*= −0.25, *p* =2.68×10^−6^) with the sample sizes, suggesting that high prediction accuracies are more likely to be achieved on a small number of subjects. Previous studies suggest that a minimum of several hundred participants are required for predictive modeling to have adequate statistical power (13, 14). Small samples may not be representative of the general population, therefore, models built using small homogeneous datasets tend to mistakenly fit sample-specific idiosyncrasies (12). Failure to account for this optimism results in the fact that neuroimaging findings are hard to generalize in practice (15). Consequently, findings derived from small sample studies often yield low reliability and should be interpreted with great caution. By contrast, the use of large datasets increases the possibility of identifying robust and generalizable brain signatures due to a higher statistical power. In this regard, predictive models based on large samples should be given more priority and emphasis, even if these studies sometimes achieve relatively lower prediction accuracies.

Importantly, the work from Sripada et *al*. is the largest predictive modeling study to date, which included 2013 subjects across 15 different sites from the Adolescent Brain Cognitive Development (ABCD) consortium (16) and predicted general cognitive ability using functional connectivity with leave-one-site-out cross-validation. The robustness of results was validated in a series of control analyses including replication in split-half analysis and in a low head-motion sample. In contrast to the leave-one-subject-out cross-validation, which has been criticized for inflated variance (12), the use of leave-one-site-out cross-validation guards against the possibility of detecting spurious brain-behavior relationships, improving our ability to identify brain signatures that will generalize.

Interestingly, a majority of the studies with large samples of subjects were performed on the publicly available data sharing initiatives, which encompass subjects spanning a range of developmental statuses and psychotic disorders. Promisingly, these data-sharing initiatives are making significant strides towards collecting large-scale neuroimaging datasets, driving the progress of brain-based biomarker discovery.

### Overview of regression-based prediction models

**Figure 2** provides an overview of regression methods that are commonly adopted in our reviewed studies. These approaches can be divided into two main classes: single-task and multi-task. Multi-task approaches jointly predict multiple phenotypic variables in a unified framework by considering the dependence relationship among behavioral measures derived from a single cognitive test (17). Only a small number of studies performed multi-task predictions, consequently, we will not present a detailed discussion here.

**Figure 2.**
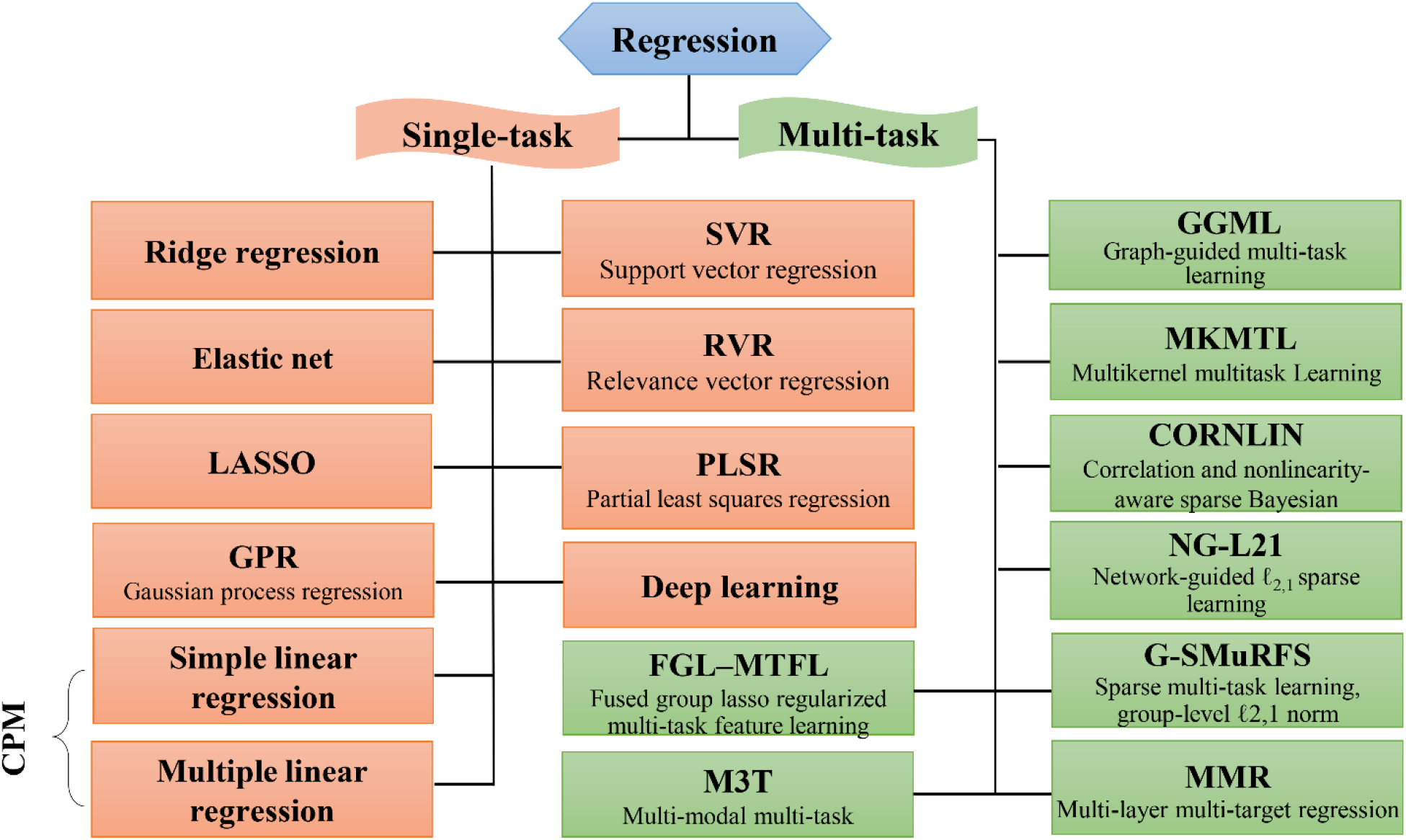
Summary of regression approaches used in our surveyed papers. Multi-task approaches jointly predict multiple clinical variables in a unified framework, while single-task methods only predict one type of cognitive score at one time. Most of the surveyed papers used linear models to reveal brain-behavior relationships. Connectome-based predictive modeling (CPM) is a recently developed data-driven approach that combines simple linear regression and feature selection together to predict individual differences in traits and behavior from connectivity data, and has been successfully employed for the prediction of multiple human behaviors.

Single-task methods predict only one type of phenotypic measure at one time. Such type of studies account for more than 90% of all surveyed papers. These methods primarily include the simple/multiple linear regression, relevance vector regression (RVR), linear support vector regression (SVR), ridge regression, LASSO, elastic net, partial least square regression (PLSR), and Gaussian process regression (GPR). Almost all of these methods belong to linear models, which are based on the hypothesis that there exists a linear relationship between brain imaging measurements and behavior scores. Although nonlinear approaches may be better suited for capturing complex brain-behavior relationships (10), the linear models could substantially reduce the possibility of overfitting and ensure good generalizability. Moreover, linear models have more interpretability, thereby allowing researchers to easily pinpoint the predictive brain regions and quantify their contribution by mapping them back to the original feature space and extracting their beta coefficients (10, 18). Here, we present some representative prediction models as follows.

#### Simple/multiple linear regression

Among all approaches, the simplest and most prevalent method for establishing brain-behavior relationship is simple/multiple linear regression (10). However, multiple linear regression requires the sample size to exceed feature dimension, and tends to overfit when the data is noisy. Given the high dimensional nature of neuroimaging data, these approaches are commonly accompanied by a feature selection step to obtain low-dimensionality representations (19). Typically, connectome-based predictive modeling (CPM) is one of such approaches that combine simple linear regression and feature selection together to predict individual differences in traits and behavior from connectivity data (3, 10, 20). **Figure 3A** shows a schematic summarizing the CPM pipeline. The CPM procedure has several advantages including straightforward interpretation, fast computation, and robust generalization, and has been successfully employed for the prediction of multiple human behaviors (20–23). **Figure 3B-C** show two example studies using CPM to predict individual differences in creativity and sustained attention.

**Figure 3.**
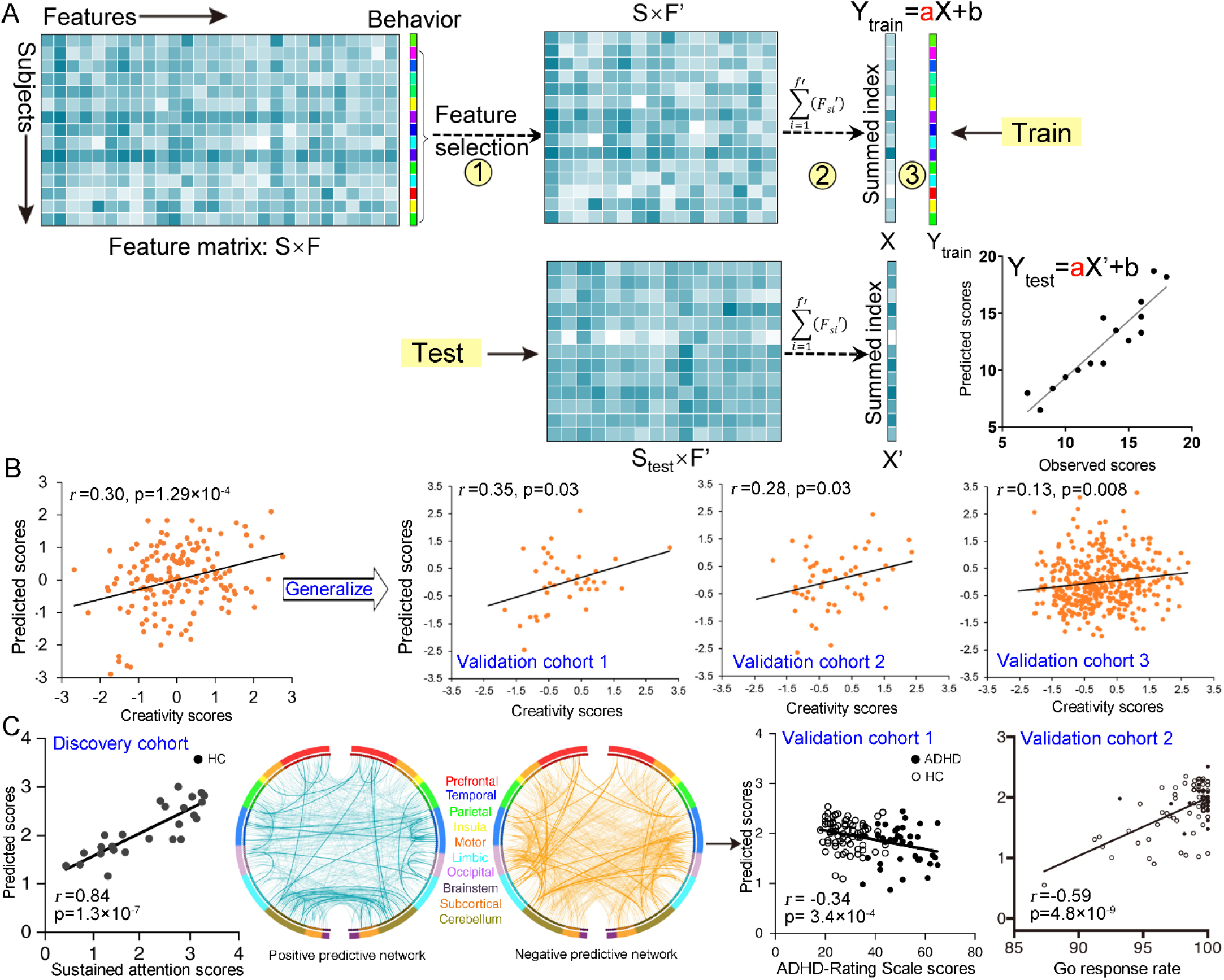
Overview of the connectome-based predictive modeling (CPM) framework and its applications in behavior prediction. (**A**) Overview of general analysis strategy for CPM procedure. Specifically, CPM is performed first by calculating the relevance of each connectivity feature to behavioral measure of interest across subjects and retaining the most significantly correlated ones under a predefined threshold. And then, a single aggregate metric named ‘network strength’ is computed by summing strength of the retained connectivity features. Afterwards, the summary statistics and behavioral scores are submitted to a simple linear regression model. By placing the procedure within a cross-validation framework, accurate estimations of behavioral scores can be obtained. (**B**) Beaty et *al*. accomplished robust prediction of individual creative ability using FCs acquired from 163 participants engaged in a classic divergent thinking task, and assessed the generalizability in three external cohorts (*n*=39, 54, and 405). (**C**) Rosenberg et *al*. accomplished robust prediction of sustained attention using CPM in a sample of 25 healthy subjects, and the identified neuromarkers were generalized to predict a clinical measure for patients with ADHD (validation cohort 1, *n*=113) and individuals’ go response rate in a stop-signal task (validation cohort 2, *n*=83). (Adapted from ref. (48), and (20, 36)).

#### LASSO

LASSO is a regularized regression method using an L1-norm penalty to reduce model complexity. LASSO shrinks most regressors’ coefficient with low predictability to zero, and retains one arbitrary feature among the correlated ones (13). This means that some important features may be absent from the final model, which may lead to problems in feature interpretation.

#### Ridge regression

The ridge regression also employs the regularization technique to impose sparsity constraints. Ridge regression includes all features in the final prediction model and assigns similar coefficient values to the correlated features. Consequently, it has a good resilience to multicollinearity (24).

#### Elastic net

Elastic net can be seen as a combination of LASSO and ridge regression. It can not only achieve a sparse model through dropping features with low predictability but can retain groups of correlated features, thereby improving interpretability and stability (25).

#### SVR

To reduce model complexity, SVR works by placing constraints to ensure only a small number of observations (support vectors) are used. SVR works with the goal of constructing a regression line that fits the data within some chosen level of error (26). Due to an easy availability of implementation tools, SVR has been widely employed for behavior predictions.

#### RVR

RVR is a sparse kernel learning regression method based on Bayesian framework (27). Like SVR, only a relatively small number of samples (relevance vectors) are used to fit the model. RVR requires no extra computation for parameter tuning, therefore, it has a good generalizability and moderate computational cost.

#### PLSR

PLSR works by representing the high-dimensional features with a small number of latent components and then using these latent components to make predictions (28, 29). Consequently, PLSR does not need a feature selection step to reduce the feature dimension, and is particularly helpful in situations where the predictors are highly collinear and high dimensional.

### Study review of individualized prediction of human behavioral measures

The regression-based machine learning models have been successfully applied in the prediction of several important behavioral aspects including cognition abilities, symptom severity, personality traits, emotion feelings and motor performance (**Figure 1C**). We now review these translational neuroimaging findings and present some example studies.

#### Cognition

As a core function of humans, cognition ubiquitously pervades one’s daily life and plays a crucial role in determining how individuals understand, learn and communicate with the world. A majority of all reviewed papers focused on predicting cognition metrics. Such investigations attempt to unravel the secret of cognition processing by examining how the brain gives rise to cognition. General intelligence, attention and reading comprehension ability constitute the top three most studied cognitive metrics (**Figure 1D**).

Prediction of cognition has been most active in general intelligence (30–32). One recent study predicted the fluid intelligence scores in a sample of 515 subjects based on functional connectivity features, and confirmed the robustness of results by controlling for an array of potential factors including cross-validation method, head motion effect, global signal regression, and brain parcellation schemes. More importantly, the prediction models could be generalized to predict three intelligence-related measure in a large independent dataset (*n*=571) (21). With respect to attention, Rosenberg et *al*. accomplished robust prediction of sustained attention using CPM in a sample of 25 healthy subjects (20), and more importantly, the identified neuromarkers were generalized to predict a series of attention-related measures like inhibition control, reading recall, and even a clinical measure for patients with attention-deficit/hyperactivity disorder (ADHD) in several independent data sets (33–36) (**Figure 3C**). Within a three-fold cross-validation framework, another study employed elastic net to decode individual differences in reading comprehension ability from grey matter volume for 507 healthy subjects, and further evaluated the generalizability in two external validation cohorts (*n*=372 and *n*=67) (37).

Apart from the aforementioned studies, predictive modelling has also been extended to achieve predictions for other cognitive metrics, including working memory (38–40), verbal learning (41), inhibition control (42, 43), processing speed (44, 45), cognitive flexibility (42, 46, 47), creativity (48–50) and spatial orientation (51) using neuroimaging features of functional or structural connectivity, grey matter volume, cortical thickness and fractional anisotropy.

#### Symptom severity

In the clinical domain, most existing criteria to assess severity of brain disorders predominantly rely on subjective judgement of the patient symptoms and self-reported history. Machine learning-based predictive modeling has been utilized to decode symptom severity or cognition dysfunction from neuroimaging data. These models can establish the quantitative relationship between symptom scores and brain changes, which can further help us track the progress of neurological diseases and better understand the pathophysiology (52). Such models have been applied to patients with a spectrum of neurological or mental health disorders, such as schizophrenia (53, 54), autism spectrum disorder (ASD) (25, 55), depression (56–58), Alzheimer’s disease (AD) (59–62), ADHD (22), Huntington’s disease (26), obsessive-compulsive disorder (63, 64), and Parkinson’s disease (65, 66). For example, based on cortical thickness measurements, Moradi et *al*. predicted the symptom severity scores for 156 subjects with ASD from four different sites (25). Another study presented a deep learning model for the prediction of clinical scores of disease severity for AD patients using structural MRI data (67). Within 10 rounds of 10-fold cross-validation, this model achieved high prediction accuracies across two independent large cohorts (*n*=669 and *n*=690).

Interestingly, models developed to predict behavioral measures for patients with brain disorders achieved significantly higher accuracies than models developed for health (p=8.12×10^−11^, Hedge’s *g*=0.73, **Figure 1E**). A potential reason may be behavioral measures show small variations in the healthy controls instead of patients, which may be too low for the algorithms to pick up on predictive features. In contrast, many clinical measures are designed to pick up differences in patients. Notably, this result should be interpreted with caution, since these studies varied in a wide range of aspects, like the phenotypic measures, prediction approaches, sample size, and cross-validation schemes.

#### Personality

Personality is a relatively stable trait consisting of a collection of behaviors, thoughts, cognitions, and emotional characteristics that evolve from biological and environmental factors (68). Personality represents an individual’s disposition that influences long-term behavioral style (24). Among all personality dimensions, traits measured using the Five-Factor Inventory, which encompasses five broad dimensions of extraversion, neuroticism, agreeableness, conscientiousness and openness to experience, have been most widely investigated. For example, within a leave-one-family-out cross-validation, Dubois et *al*. predicted the scores of openness and a personality factor generated from the principal component analysis using elastic net in a large sample of 884 adults (24). Another study, based on nine meta-analytically derived functional networks and RVR, demonstrated the predictability of extraversion, neuroticism, agreeableness and openness within a 10-fold cross-validation in a sample of 420 subjects, and assessed the generalizability of findings in a replication sample of 310 subjects (69). Moreover, this study suggested that the functional networks linked to each personality dimension were gender-specific. Other personality traits being investigated primarily include temperament (70, 71), narcissism (72), and dispositional worry (27).

#### Emotion

Only a limited number of studies have attempted to identify biomarkers for emotion related behaviors using predictive modeling, which have largely deepened our understanding of the underlying neurobiological substrates. These emotion measures primarily include individuals’ feeling of loneness (73), anxiety (74–76), fear of pain (77), intensity of interoception (78), as well as constructs associated with social relationships like propensity to trust (79) and deception (80).

Apart from the above mentioned phenotypic aspects, there are also some other behavioral domains being investigated (81, 82). A remarkable example is the study that identified an fMRI-based brain signature of pain intensity, and showed its generalizability in multiple independent datasets (81). Due to word limits, we will not describe these studies in detail.

## Challenges and future directions

The studies surveyed above highlight the potential of using regression-based predictive modeling to identify neuromarkers and characterize the neurodiversity of the human brain in both health and disease. Despite such success, some issues should be mentioned and lots of work remains to be done.

### Prediction using Multimodal data

Recently, collecting multimodal brain data from the same subject has become a common practice (41, 83, 84). Integrating multimodal data could effectively capitalize on the strength of each imaging modality and provide a comprehensive view into the brain (85). As such, in the surveyed papers, studies using multimodal data to predict individuals’ behaviors achieved significantly higher accuracies than those using unimodal data (p=1.45×10^−6^, Hedge’s *g*=0.72, **Figure 1F**). A recent study predicted the intelligence quotient scores using resting-state functional connectivity and grey matter cortical thickness, and found that integrating multimodal data improved prediction accuracy (86). More importantly, this study suggested that these two types of neuroimaging features provided unique evidence of the neurobiological correlates of intelligence from distinct perspectives (**Figure 4A**).

**Figure 4.**
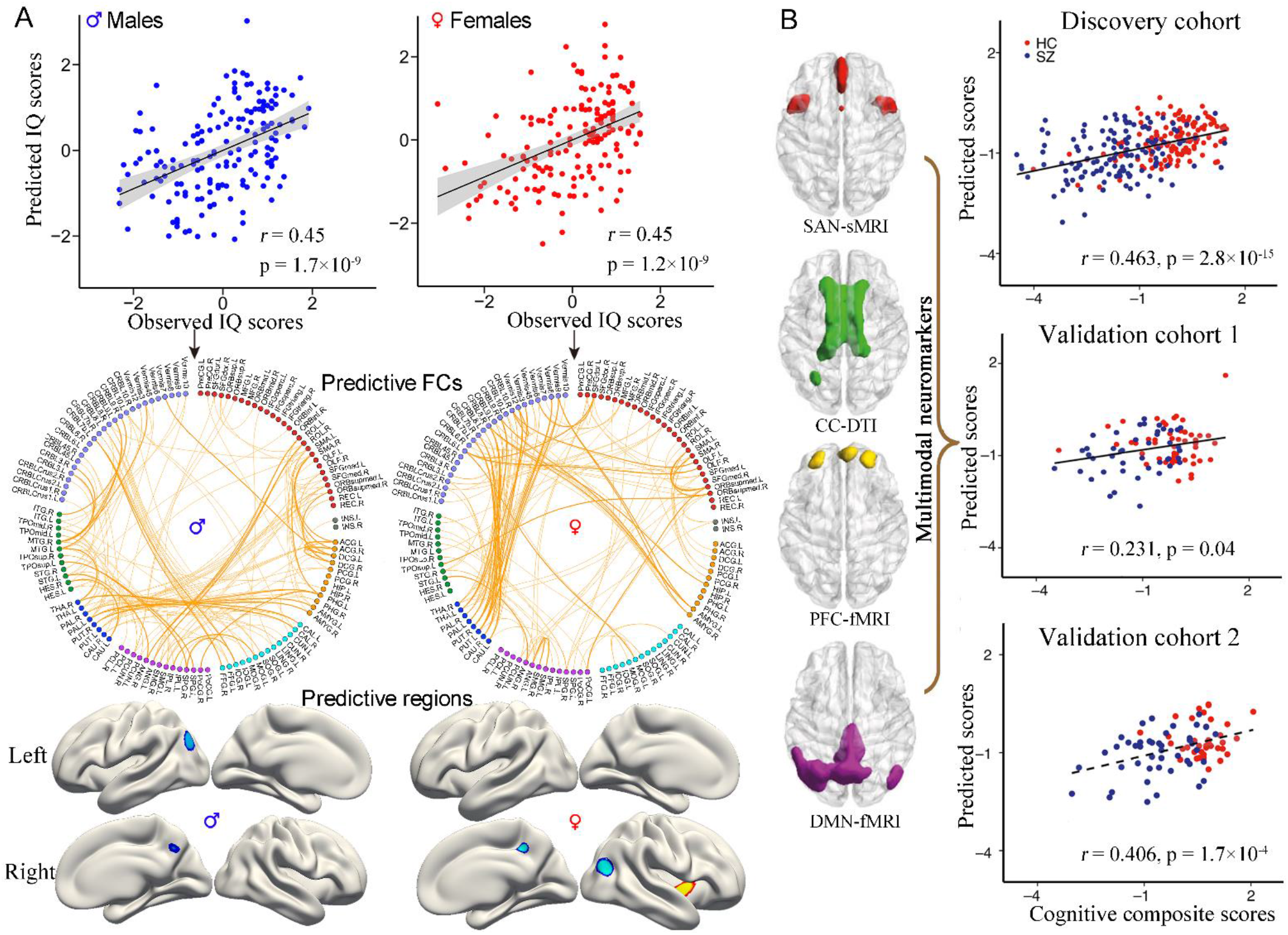
Using multimodal data to predict cognition promotes the biomarker identification. (**A**) Jiang et al. achieved an improved prediction performance of intelligence scores by integrating FCs and cortical thickness. More importantly, the study suggested that these two types of neuroimaging features provided unique evidence of the neurobiological correlates of intelligence from distinct perspective. Specifically, prediction with cortical thickness explored more gender difference in the lateralization of predictive brain regions, while prediction with FCs detected more gender difference in the specification of contributing functional networks. (**B**) Sui et *al*. predicted the cognitive composite scores for subjects from three independent cohorts based on neuromarkers derived from a supervised multimodal fusion approach (Adapted from ref. (86), and (87)).

However, the multimodal prediction was primarily achieved by simply concatenating brain features from different modalities horizontally into a single, combined feature space and submitting them into a regression model, thereby not allowing for a full use of the joint information among modalities. Multimodal fusion can uncover cross-information beyond what can be detected by each single modality (85). For example, Qi *et al.* proposed a supervised multimodal fusion approach named MCCAR+jICA, which can detect co-varying multimodal imaging patterns (84). In a three-way fusion simulation, the method identified four multimodal brain signatures located in the salience network, central executive network and default mode network, which were successfully used to predict multi-domain cognition in two independent cohorts (87) (**Figure 4B**). The use of brain signatures derived from multimodal fusion approaches to predict cognition can promote the identification of neuromarkers and inform our understanding of how functionally and structurally connected brain systems contributed to cognitive function.

### Validating biomarkers in independent data sets

Many existing studies detecting neuromarkers predominately focus on revealing new findings for a cascade of novel behavioral measures. However, additional effort should also be devoted to testing the generalizability and reproducibility of the constructed predictive models (88), especially for those developed on dataset with an insufficient sample size. Considering a potentially inflated false-positive rate due to the low statistical power, concerns have been raised about the reliability and reproducibility of neuroimaging findings (89, 90). However, very few studies have tested brain models on external validation/replication datasets due to generalization failure or lack of additional independent data. One potential solution to this issue is to encourage individual projects to share their neuroimaging data so that researchers can leverage the publicly available data. The shared big-data sources can serve as benchmark data sets, allowing people to establish reproducible imaging-based biomarkers (91). Moreover, the neuroimaging community should raise awareness that replication and validation are as important as novelty (89).

### Using participants’ performance in a task to assess a trait

In practice, assessments of behavioral measures can be based either on subjective report or on objective behavioral recordings. The self-reported measures are usually derived from subjects’ ratings of a conventional scale according to their personal experience or judgement on a specific item. This means that the measured scores rely largely on individuals’ subjective appraisal of the trait. Consequently, such measures are likely to be influenced by unknown factors and are not stable. In contrast, behavioral measures assessed using participants’ performance in a task where participants are incited to reach a high score (92), objectively reflect ones’ inherent traits and relate to more reliability, placing an upper limit on the maximum detectable effect size (93). Some studies have reported that objective variables can be better predicted than subjective ratings (94, 95). In addition, it has also been suggested that cognitive tasks amplified trait-relevant individual differences in functional connectivity patterns and integrating different tasks into a single predictive model did improve cognition prediction (21, 96, 97). Therefore, we encourage future studies to use task performance to measure individuals’ behavioral scores and combining multiple task connectomes to investigate brain-behavior relationships.

### Applying deep learning methods

Recent studies have started to use deep learning methods to predict individual differences in behavioral phenotypes and brain maturity. Based on neural networks, deep learning serves as an extension of traditional machine learning methods (98). Deep learning can automatically learn a higher level of abstract representation through the application of consecutive nonlinear transformations to raw input data in “hidden” neural network layers, making this approach ideally suited to detecting complex, scattered and subtle brain patterns (99). One advantage of deep learning is that it can remove the reliance on time-consuming pre-processing and prior feature selection, avoiding the related model-dependent decisions (100). However, compared to other fields, the application of deep learning methods to predictive modeling in neuroimaging is relatively modest. There are two main reasons, one being the lack of interpretability and the other being the need for extensive amounts of data. Specifically, one study suggested that, lacking large quantity of data, deep learning models may not outperform classical machine learning approaches in cognition predictions (101). Therefore, many efforts are required to overcome these drawbacks before the full potential of deep learning in behavior prediction is explored.

### Developing longitudinally predictive models

The vast majority of predictive studies have focused on cross-sectional predictions, that is, the MRI data and the behavioral measure of interest are acquired at the same time or within a short time interval. Although there is evidence showing that certain brain imaging features are unique and stable over months to years (102), to what degree these features will show predictability for behavioral phenotypes consistently across time still remains largely unexplored. To benefit educational or healthy practices, longitudinal predictive models should be developed to predict long-term outcomes using baseline neuroimaging data (2, 11). Biomarkers derived from such applications commonly relate to more biological significance and clinical utility, since individuals can benefit from early invention and treatment.

## Conclusions

In this study, we reviewed recent advances in the field of predictive modeling that employs regression-based machine learning approaches to decode individual differences in behavioral phenotypes from brain imaging data. Although this burgeoning field is still immature with many issues being solved and not quite ready for integration into clinical use, we are optimistic about the development of brain models that can be eventually integrated into clinical applications as this area matures.

## Acknowledgments

This work is supported in part by the National Institute of Health (1R01MH117107, R01EB020407, P20GM103472, P30GM122734) and the National Science Foundation (1539067), the China Natural Science Foundation (61773380), the Strategic Priority Research Program of the Chinese Academy of Sciences (No. XDB32040100), Brain Science and Brain-inspired Technology Plan of Beijing City (Z181100001518005),

The authors declare no biomedical financial interests or potential conflicts of interest.

## Supplementary File

**Table S1.**
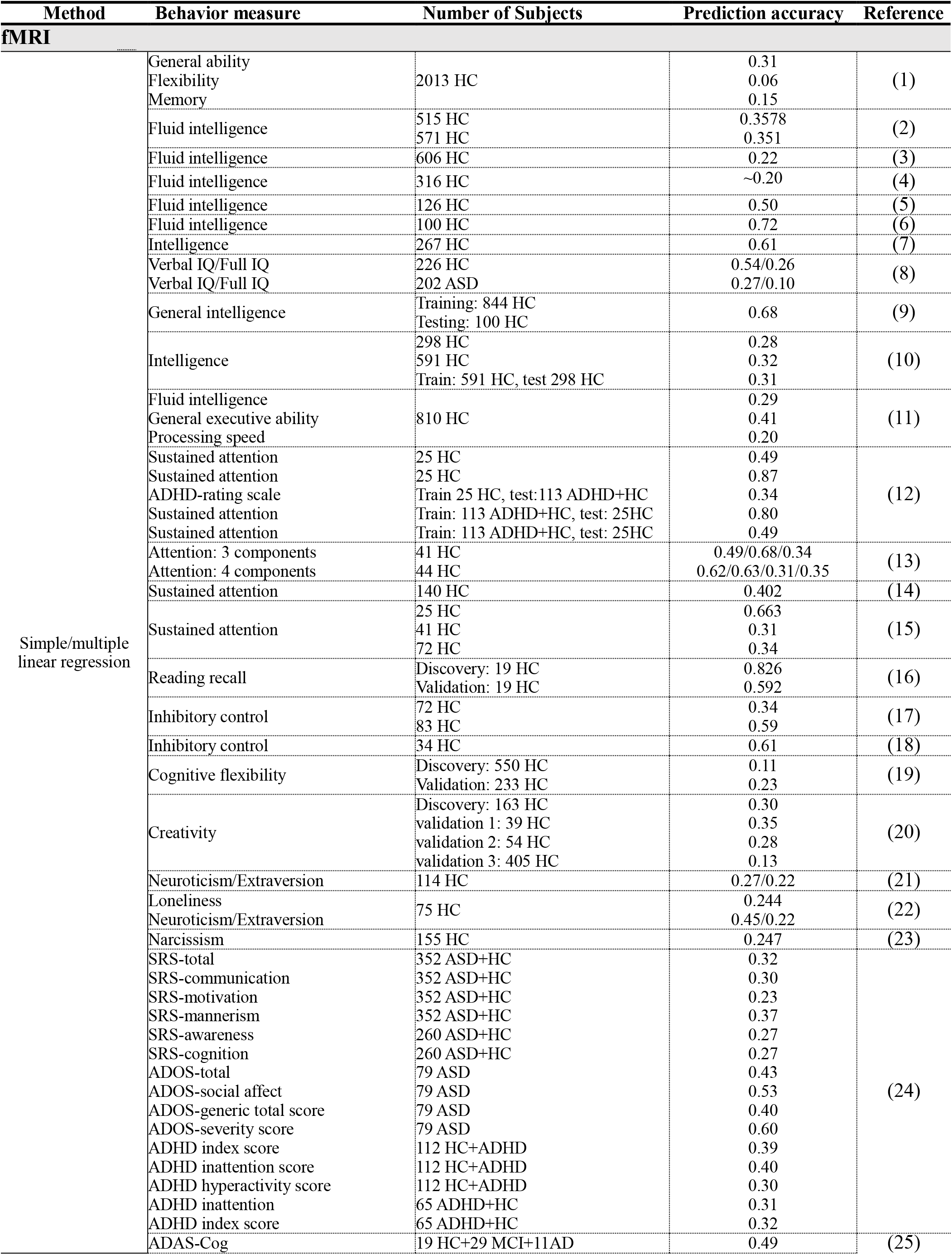
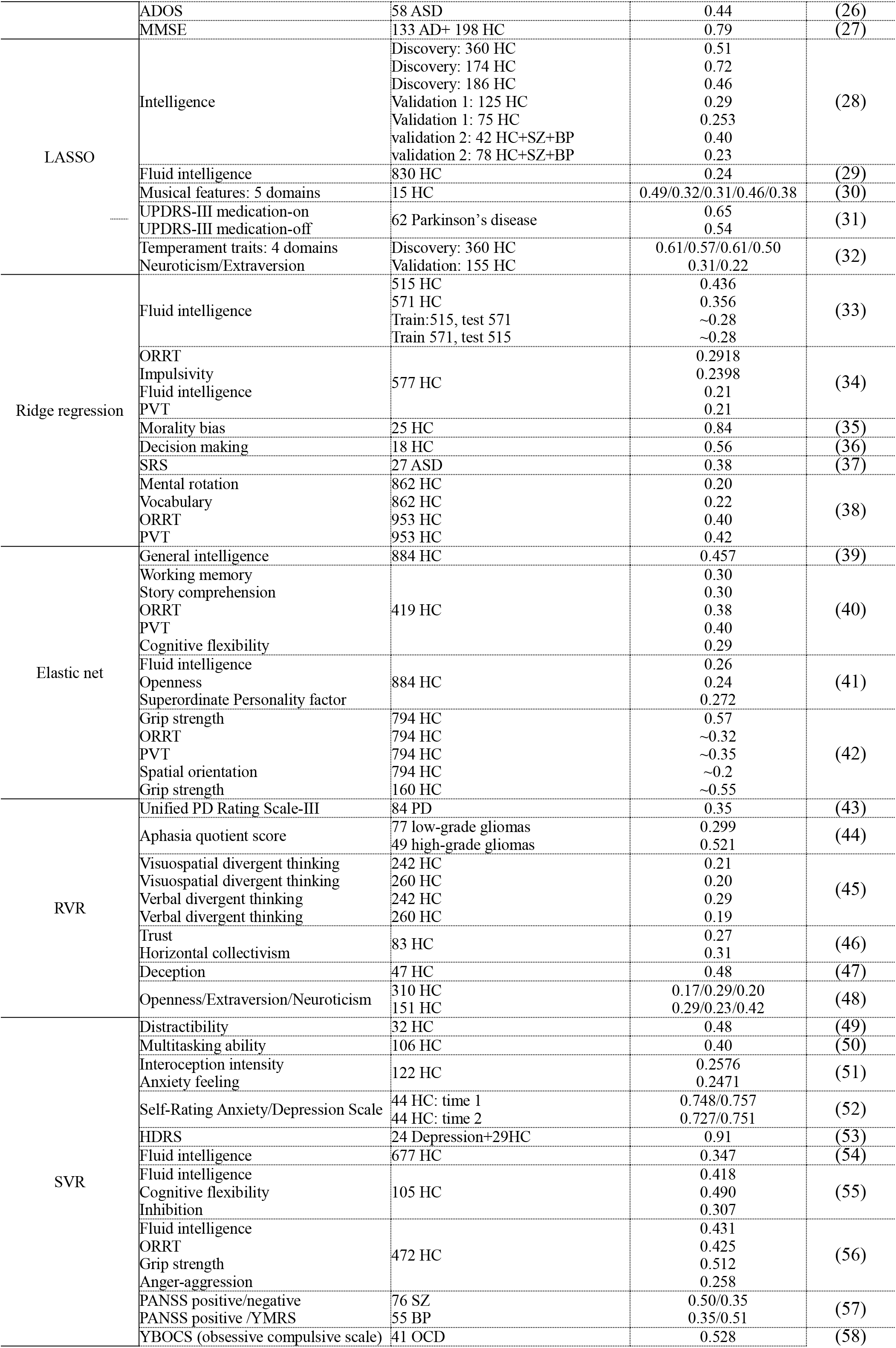
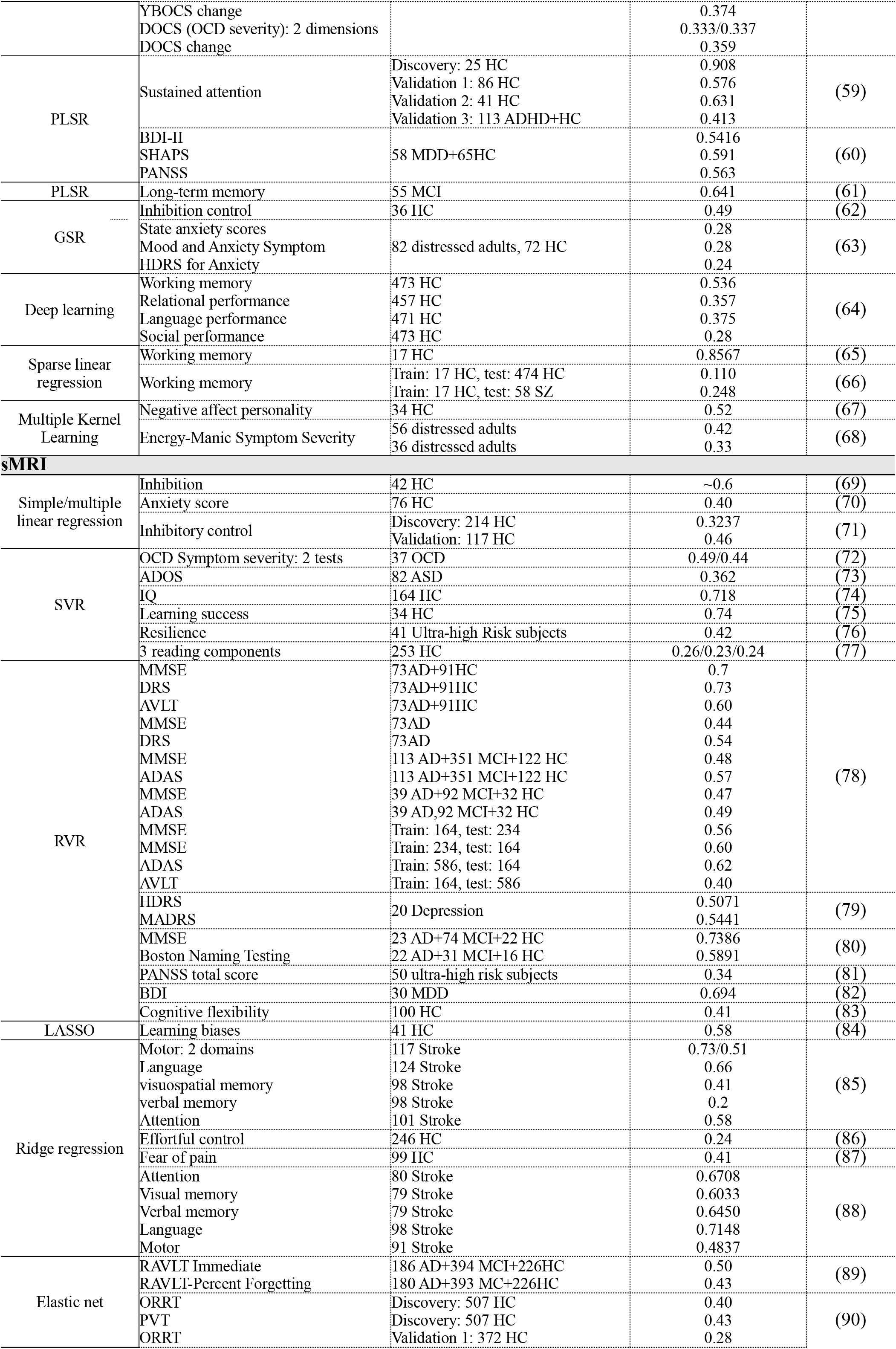
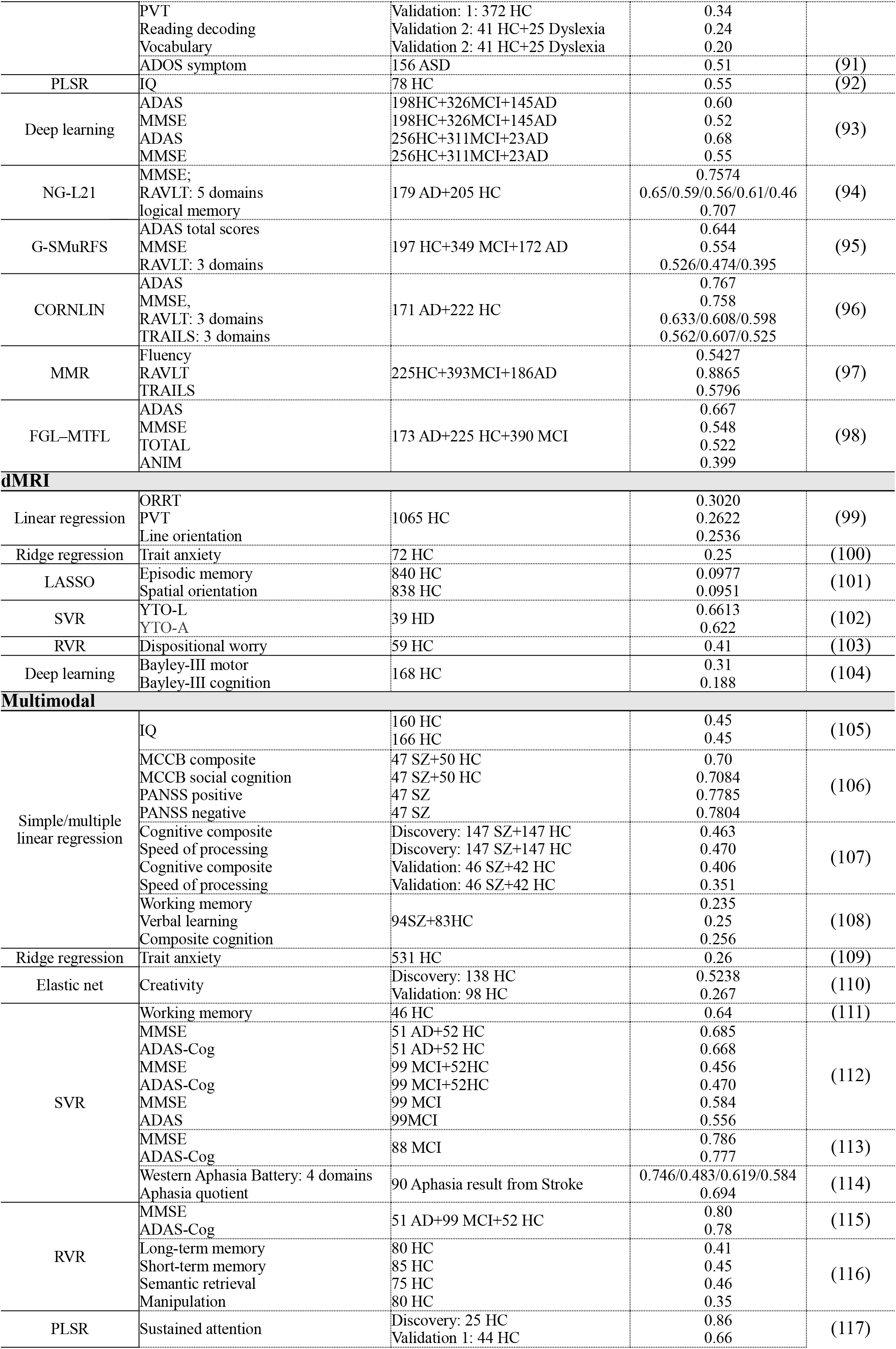
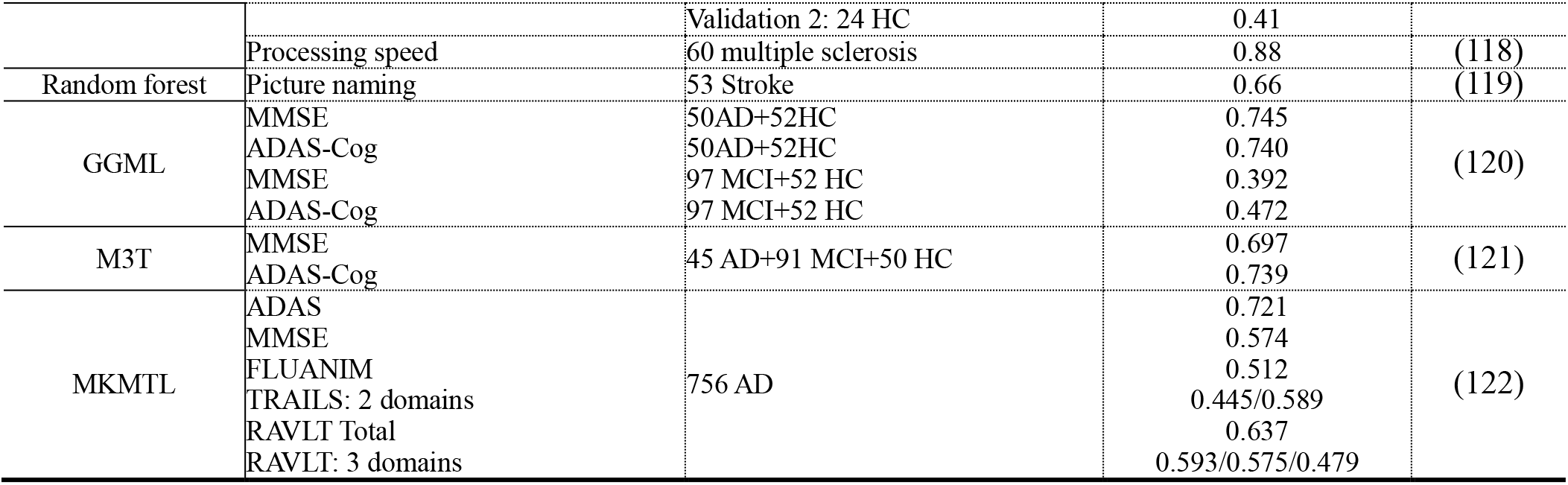
Summary of 122 regression-based prediction studies.

